# Persistence without turnover: the RhoG G12E mutant highlights the role of nucleotide cycling in RhoG signaling

**DOI:** 10.64898/2026.03.25.713116

**Authors:** Shah Wajed, Yann Ferrandez, Mahel Zeghouf, Rasta Ghasemi, Cylia Puertas, Agata Nawrotek, Franck Gesbert

## Abstract

Small GTPases of the Ras superfamily act as molecular switches cycling between GDP-bound inactive and GTP-bound active states. Their signaling output depends not only on GTP loading but on continuous nucleotide cycling controlled by GEFs and GAPs. While Gly12 substitutions in Ras typically confer constitutive activation, the effects of equivalent mutations in Rho-family GTPases remain poorly defined. Here, we characterize a ClinVar-reported RhoG^G12E^ variant. Quantitative kinetic studies show that RhoG^G12E^ has strongly impaired intrinsic and GAP-stimulated GTP hydrolysis but it can still be activated by GEFs. Effector binding is preserved *in vitro* and RhoG^G12E^ accumulates in a GTP-bound state in cells. Yet, RhoG^G12E^ induces enhanced spreading, increased focal adhesions, and reduced collective migration – phenotypes consistent with previously reported consequences of reduced RhoG signaling rather than hyperactivation. These findings indicate that accumulation of the GTP-bound form of small GTPases does not necessarily translate into productive output, supporting a model in which impaired nucleotide cycling compromises RhoG-dependent cellular behaviors.

## Introduction

Small GTPases of the Ras superfamily operate as nucleotide-dependent molecular switches that cycle between inactive GDP-bound and active GTP-bound states. This cycle is spatially organized by GEFs (Guanine nucleotide Exchange Factors) that promote activation at membranes in response to upstream signals, enabling effector engagement, whereas GAPs (GTPase-Activating Proteins) accelerate GTP hydrolysis to terminate signaling once the stimulus has been decoded [1]. Glycine 12 (Gly12) is a highly conserved residue located in the P-loop (phosphate-binding loop) of Ras superfamily GTPases, where it directly contributes to positioning the β- and γ-phosphates of the bound nucleotide. In Ras, substitution of the conserved Gly12 leads to a constitutively active form thereby sustaining downstream Ras signaling pathways promoting oncogenic transformation and uncontrolled proliferation [2],[3]. Given that Ras and Rho proteins both operate within this evolutionarily conserved nucleotide-switch paradigm and are governed by analogous regulatory mechanisms, their functional analysis has often employed analogous mutational strategies. This paradigm rests on the assumption that locking a small GTPase in the GTP-bound state is sufficient to maintain signaling in a constitutively active output.

Whether this interpretation can be directly transposed to Rho-family GTPases remains unclear. In contrast to many Ras outputs, numerous Rho-dependent processes—such as actin remodeling, adhesion turnover, and directed migration—require rapid and spatially confined nucleotide cycling [4], [5]. Under such conditions, persistent GTP loading without efficient turnover may not faithfully reproduce physiological activation and may instead perturb signaling dynamics. The recent identification of mutations affecting this conserved Gly residue in Rho GTPases, associated with pathological conditions, now compels a direct evaluation of whether the constitutive activation paradigm established for Ras can be transposed to the Rho family [6].

In the present study, we identified in a public archive of human genetic variants (ClinVar database) a RhoG^G12E^ variant (ClinVar ID: NM_001665.4(RHOG):c.35G>A (p.Gly12Glu)) representing, to our knowledge, the only described mutant of RhoG on position 12 reported in patients in the context of immune-related phenotypes. RhoG is a member of the Rac subfamily of Rho GTPases and plays a key role in actin remodeling, membrane protrusion, and cell migration, in particular through activation of the ELMO–DOCK signaling axis [7],[8],[9]. It has been reported that siRNA mediated knock down of RhoG leads to a delay in EGF-induced migration of HeLa cells in wound healing assays and RhoG-depleted cells show defective migration and excessive adhesion [10],[11].

However, the biochemical and cellular consequences of a Gly12 substitution in RhoG have never been examined.

At a structural level, Gly12 provides steric flexibility within the P-loop that supports proper Mg^2+^ coordination and alignment of catalytic elements required for efficient hydrolysis[12]. Replacement by glutamate (G12E) in RhoG introduces a negatively charged side chain in this constrained environment, potentially perturbing the geometry of the nucleotide-binding pocket (**Supp.Fig.1**). Such a substitution is expected to impair GAP-stimulated hydrolysis and may alter the kinetic balance between activation and inactivation without necessarily abolishing nucleotide binding or effector recognition.

Here, we combine quantitative biochemical assays with cell-based analyses to dissect the consequences of the G12E mutation in RhoG. We show that RhoG^G12E^ loses its intrinsic and GAP-stimulated GTP hydrolysis, resulting in a severe imbalance in regulatory control. Despite this, RhoG^G12E^ retains effector-binding capacity and accumulates in an active state in cells. Surprisingly, at the cellular level, expression of RhoG^G12E^ drives pronounced actin cytoskeleton reorganization, increased cell spreading, enhanced focal adhesion formation, and reduced migratory capacity that resemble RhoG depletion rather than hyperactivation. These findings indicate that persistence in the GTP-bound state does not equate to productive signaling and highlight the importance of nucleotide cycling, rather than static GTP loading, for RhoG-dependent cellular functions.

## Results

### RhoG G12E mutation results in reduced EDTA-stimulated nucleotide exchange

Mutations at position 12 of small GTPases are known to perturb the GTPase cycle, but the specific biochemical consequences vary widely across family members. We first sought to determine whether G12E alters nucleotide exchange, i.e., the ability of RhoG to release GDP and bind GTP. Spontaneous nucleotide exchange was monitored using BODIPY-GDP-loaded RhoG proteins. Under these conditions, wild-type RhoG exhibited a slow but measurable GDP-to-GTP exchange rate (k_obs_ = 2.51±0.31 × 10^−4^ s^−1^), and RhoG^G12E^ exchanged at a comparable rate within experimental variability (k_obs_ = 3.14±1.19 × 10^−4^ s^−1^) (**Fig. 1A**). As nucleotide exchange rates were indistinguishable in the presence of Mg^2+^, we assume that G12E mutation does not destabilize the Mg^2+^-bound nucleotide complex and does not promote spontaneous nucleotide release. We therefore investigated whether the mutation affects the conformational flexibility of the P-loop/switch region that enables nucleotide release. Chelation of Mg^2+^ with EDTA removes the stabilizing ion but still requires the protein to undergo the structural transition necessary for nucleotide dissociation. As expected, EDTA-stimulated exchange accelerated nucleotide release for both proteins, however, under this condition, a clear kinetic difference was observed : RhoG^WT^ displayed twofold faster exchange rate (k_obs_ = 7.43±0.14 × 10^−3^ s^−1^), RhoG^G12E^ (k_obs_ = 3.87±0.07 × 10^−3^ s^−1^)(**Fig. 1B**). To further probe whether this effect reflected a general perturbation of nucleotide binding, we measured the reverse reaction by monitoring BODIPY-GTP-to-GDP exchange. Here again, RhoG^WT^ exhibited a faster exchange rate (k_obs_ = 6.32±1.09 × 10^−3^ s^−1^) compared to RhoG^G12E^ (k_obs_ = 3.29±0.73 × 10^−3^ s^−1^) (**Fig. 1C**). Importantly, because Mg^2+^ is chelated in both reactions, the persistence of a kinetic difference indicates that the rate-limiting barrier for nucleotide release is not Mg^2+^ removal itself. Rather, these data suggest that the G12E substitution affects a subsequent conformational transition required for nucleotide exit, consistent with altered dynamics of the P-loop/switch region. Consistent with this interpretation, apparent nucleotide affinity measured through a range of GTP concentrations remained similar between the two proteins (**Fig. 1D**), arguing against a primary defect in equilibrium nucleotide binding : RhoG^WT^ (*K*_*d*_^*app*^ = 0.19±0.01× 10^−6^ M) and RhoG^G12E^ (*K*_*d*_^*app*^ = 0.18±0.006 × 10^−6^ M) (**Fig. 1E**). Thus, the slower exchange observed for G12E under EDTA conditions points to altered switch dynamics.

**Figure 1.**
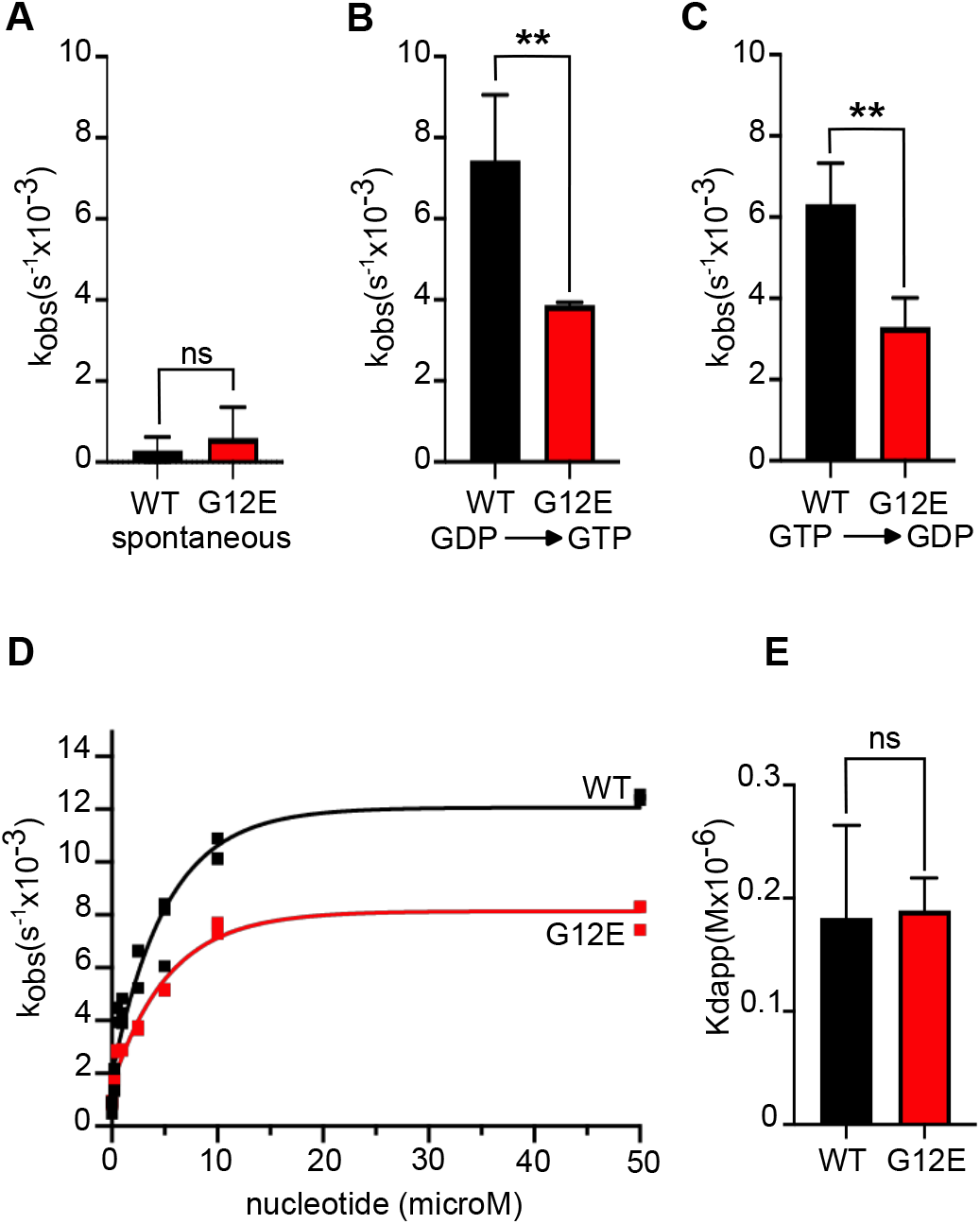
Effect of the G12E mutation on nucleotide exchange. **(A)** Intrinsic spontaneous nucleotide exchange kinetics of RhoG^WT^ and RhoG^G12E^ monitored by BODIPY fluorescence decay over time during BODIPY GDP to GTP exchange. **(B)** EDTA-induced nucleotide exchange rates monitored by BODIPY fluorescence decay over time during BODIPY GDP to GTP exchange. **(C)** EDTA-induced nucleotide exchange rates monitored by BODIPY fluorescence decay over time during BODIPY GTP to GDP exchange. **(D)** GTP titration of RhoG^WT^ and RhoG^G12E^ in the presence of EDTA (20 mM). Apparent rate constants (k_obs_) were determined over a range of GTP concentrations (0–50 µM). Each data point represents the mean of three independent experiments. **(E)** EC_50_ values derived from the curves in panel (D) were obtained by regression analysis and are reported as Kd_app_.

### G12E mutation reduces yet preserves RhoG activation by its GEFs

The EDTA condition highlights a kinetic barrier to nucleotide release in RhoG^G12E^, but physiological exchange is catalyzed by GEFs rather than by Mg^2+^ chelation. We therefore tested whether the mutation also limits catalyzed activation by measuring nucleotide exchange in the presence of the DHPH domains of PREX1 and TRIO. The activation of RhoG^WT^ and RhoG^G12E^ was assessed through measurement of BODIPY-GDP-to-GTP exchange rates across a range of GEF concentrations (**Supp Fig. 2**).

Both GEFs activated RhoG^G12E^ ~1.5–2-fold less efficiently than wild-type RhoG **(Fig.2)**. For PREX1, the catalytic efficiency on RhoG^WT^ was approximately 1.5 times faster (k_cat_/K_M_ **=** 2.91±0.20 × 10^4^ M^−1^ s^−1^) than on RhoG^G12E^ (k_cat_/K_M_ **=** 1.36±0.22 × 10^4^ M^−1^ s^−1^) (**Fig. 2A**). Similarly, TRIO-mediated activation of RhoG^WT^ was nearly twofold higher exchange rate (k_cat_/K_M_ = 2.22 ± 0.25 × 10^4^ M^−1^ s^−1^) than activation of RhoG^G12E^ (k_cat_/K_M_ **=** 1.12± 0.18 × 10^4^ M^−1^ s^−1^) (**Fig. 2B**). These results indicate that the G12E mutation affects GEF-mediated activation but does not abolish productive GEF–RhoG interactions.

**Figure 2.**
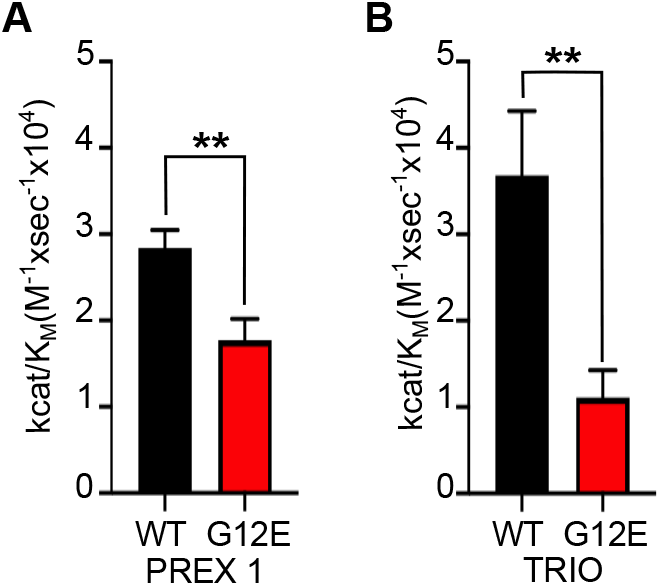
G12E mutation in RhoG decreases GEF activities of PREX and TRIO. **(A)** Catalytic efficiency of the DHPH domains of PREX1 toward RhoG^WT^ and RhoG^G12E^. **(B)** Catalytic efficiency of the DHPH domains of Trio toward RhoG^WT^ and RhoG^G12E^. The catalytic efficiency was assessed by nucleotide exchange assays monitored by BODIPY fluorescence decay over time during BODIPY GDP to GTP exchange. Apparent rate constants (k_obs_) were determined using exponential decay fit; each k_obs_ value is a mean from three independent experiments. k_cat_/K_M_ values were determined from k_obs_ obtained over a range of GEF concentrations (0–150 nM for Trio and 0–100 nM for PREX1).

### G12E mutation inhibits intrinsic GTPase activity

We asked next if the G12E mutation affects how efficiently RhoG returns to the GDP-bound state. We therefore measured first intrinsic GTP hydrolysis. A fluorescently labeled bacterial phosphate-binding sensor was used to measure the kinetics of inorganic phosphate (Pi) release [13, 14](**Fig. 3A**). Wild-type RhoG displayed readily detectable intrinsic GTPase activity (k_obs_ = 3.89±0.02 × 10^−3^ s^−1^). In contrast, the intrinsic hydrolysis activity was decreased by approximately 35-fold (k_obs_ = 0.11±0.002 × 10^−3^ s^−1^) with RhoG^G12E^ (**Fig. 3B**). These data establish that the G12E substitution strongly compromises the intrinsic hydrolysis step of the RhoG cycle, creating a pronounced kinetic bottleneck at the level of GTP hydrolysis.

**Figure 3.**
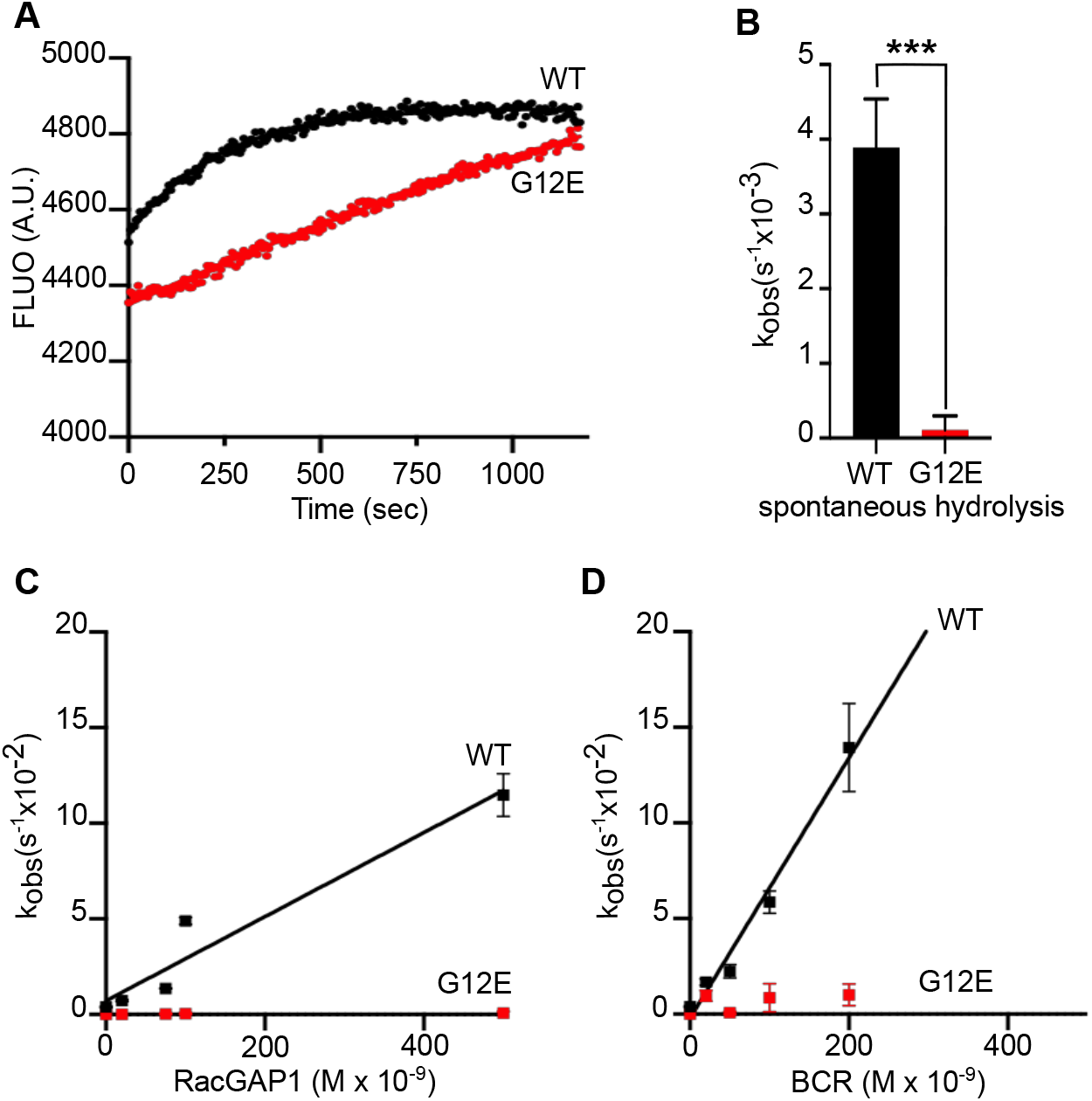
RhoG^G12E^ displays reduced spontaneous GTPase activity and decreased GAP-stimulated hydrolysis. **(A)** Example of raw curves corresponding to measured inorganic phosphate release after GTP hydrolysis over time. **(B)** Apparent rate constants corresponding to intrinsic GTPase activity for RhoG^WT^ and RhoG^G12E^. **(C)** Catalytic efficiency of RacGAP1 (GAP domain) toward RhoG^WT^ and RhoG^G12E^. **(D)** Catalytic efficiency of BCR (GAP domain) toward RhoG^WT^ and RhoG^G12E^. Apparent rate constants (k_obs_) were determined using exponential fit, each k_obs_ is a mean from three independent experiments. k_cat_/K_M_ values for RhoG^WT^ were determined from k_obs_ obtained over a range of GAP concentrations ranging from 20–200 nM.

### G12E mutation nearly abolishes GAP-stimulated hydrolysis

Intrinsic hydrolysis is slow for most small GTPases and is normally accelerated by GAPs. The strong reduction in intrinsic activity therefore raised a more decisive question: can GAPs catalyze hydrolysis of RhoG^G12E^ ? Using RacGAP1 and BCR as RhoG-active GAPs, we quantified GAP-stimulated GTP hydrolysis. The GAP rates were measured over a range of RacGAP1 or BCR concentrations, from which k_cat_/K_M_ values were determined. RacGAP1 efficiently stimulated GTP hydrolysis in wild-type RhoG (k_cat_/K_M_ = 2.16±0.03 × 10^5^ M^−1^ s^−1^), whereas activity toward RhoG^G12E^ was in the range of intrinsic activity under identical conditions (**Fig. 3C)**. Similarly, BCR exhibited strong GAP activity toward RhoG^WT^ (k_cat_/K_M_ **=** 6.80±0.05 × 10^5^ M^−1^ s^−1^), but was largely ineffective toward RhoG^G12E^ (**Fig. 3D)**. The magnitude of this defect suggests that GAP-mediated catalysis is severely impaired. Therefore, unlike GEF-mediated activation, which remains functional, GAP-stimulated hydrolysis is nearly abolished, leading to a profound kinetic imbalance.

### RhoG G12E mutation results in increased effector binding in cells

The near-complete loss of GAP activity predicts that RhoG^G12E^ accumulates in a GTP-bound state. We further investigated whether it is an effector-binding competent state. To determine whether the altered biochemical cycle of RhoG^G12E^ affects downstream signaling, we examined its interaction with the effector ELMO1 [7],[15]. *In vitro* binding assays, performed using purified RhoG protein variants and GST–ELMO1 as a bait, showed that GTP-loaded RhoG^WT^ and RhoG^G12E^ bound GST–ELMO1 with comparable efficiency, while GDP-loaded forms, as expected, bound in a weaker manner (**Fig. 4A**). These results indicate that the G12E substitution does not intrinsically alter ELMO1 recognition. The ELMO1-RhoG interaction was thus further examined to characterize the pools of active forms of RhoG variants in cells. Using a lentivirus-based transduction approach, we generated HeLa cell lines stably expressing either RhoG^G12E^ or RhoG^WT^. Immunoblot analysis confirmed that both cell lines expressed comparable levels of RhoG, with expression similarly elevated relative to parental cells (**Supp Fig. 3A and B**). Purified GST-ELMO1 was used to perform pull-down experiments of active RhoG from cell lysates. As controls, when cell extracts were treated to load GTPases with GDP or GTP (**Fig. 4B**, lanes 1-3 and lanes 4-6 respectively) we observed a significantly greater binding of RhoG in the GTP condition. Using untreated lysate, pull-down assays revealed more RhoG recovered from cells expressing RhoG^G12E^ than from cells expressing RhoG^WT^ (**Fig. 4B**), while equivalent amounts of RhoG variants were expressed in both inputs (**Fig. 4C**). Thus, increased effector binding reflects accumulation in a GTP-bound state rather than enhanced affinity.

**Figure 4.**
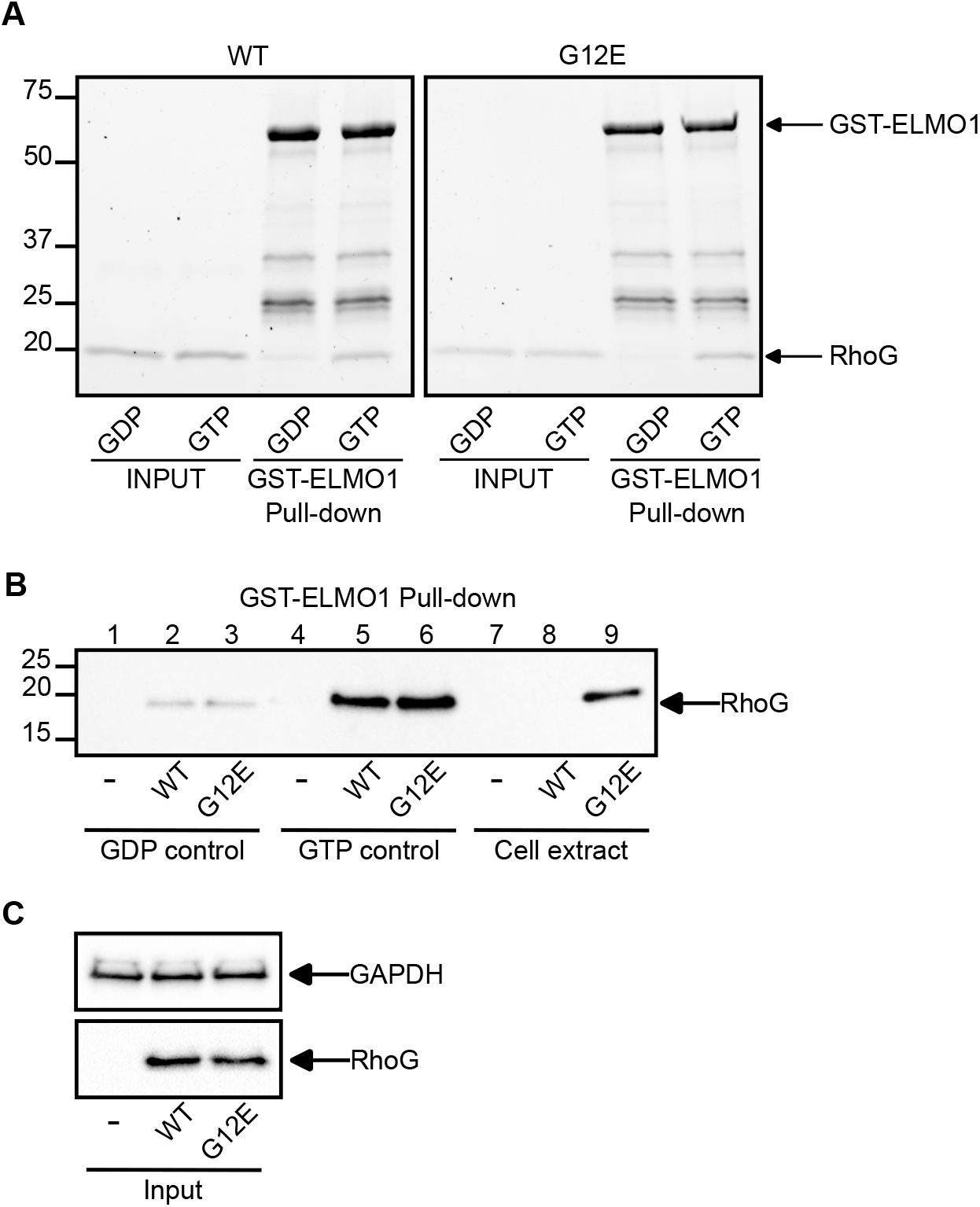
RhoG^G12E^ expression leads to the accumulation of active RhoG in cells. **(A)** Coomassie blue stained SDS-PAGE gel from pull-down assay with purified RhoG^WT^ and RhoG^G12E^ proteins in controlled GDP or GTP-bound states. The GTP-loaded forms of both proteins interact strongly with GST-ELMO1-coated beads. The results were reproduced in three independent experiments. **(B)** GST-ELMO1 mediated pull-down of active RhoG in HeLa parental, RhoG^WT^, and RhoG^G12E^ expressing cells. Lanes: GDP-treated negative control (1–3), GTP-treated positive control (4–6), and cell extract only (7– 9). **(C)** Input control showing total RhoG expression levels in crude lysates of parental, RhoG^WT^, and RhoG^G12E^ cells, with GAPDH loading control.

### Expression of RhoG^G12E^ variant triggers cell spreading and modification of cell morphology

We next asked whether accumulation of active RhoG^G12E^ translates into functional cellular phenotypes. To this end, the established cell lines described earlier were seeded at a similar confluency and treated for scanning electron microscopy or for immunofluorescence studies (**Material and methods**).

Scanning electron microscopy shows that RhoG^G12E^-expressing cells are flatter and more spread out compared to RhoG^WT^-expressing cells, which resembled parental control HeLa cells more closely (**Fig. 5A.a and b, Supp. Fig. 4A-B**). We further investigated these morphological changes in cells by imaging filamentous actin using confocal microscopy (**Fig. 5A.c and d**). RhoG^G12E^-expressing cells displayed a pronounced reorganization of the actin cytoskeleton, characterized by extensive cortical actin organization and an increase in cell protrusions that may resemble enlarged lamellipodia structures. Quantification of cell area confirmed that RhoG^G12E^-expressing cells were significantly larger than either RhoG^WT^ or parental cells (**Fig. 5B and Supp. Fig. 4. A-B**), consistent with altered cytoskeletal regulation.

**Figure 5:**
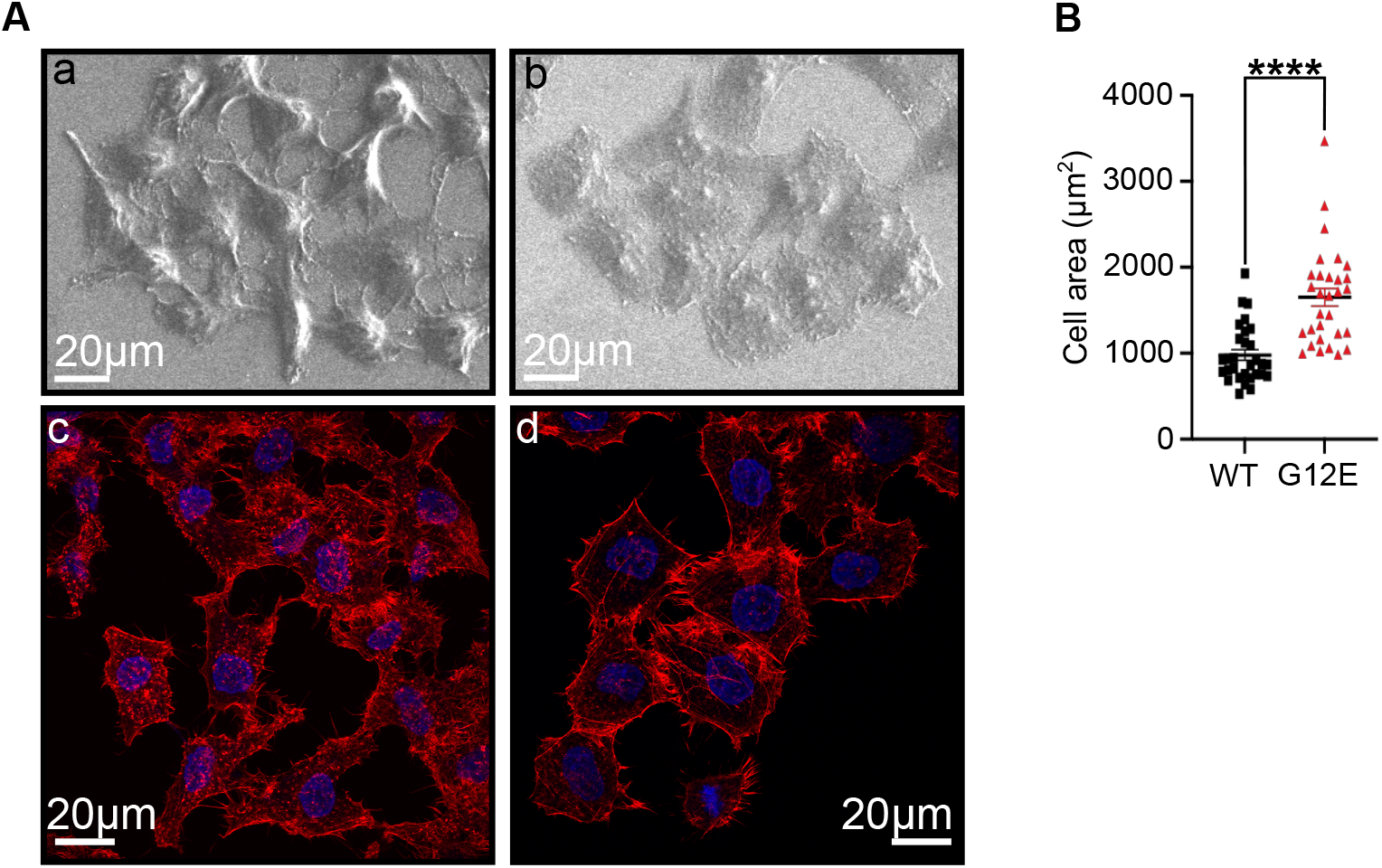
RhoG^G12E^ expression leads to increased cell-spreading. **(A)** Scanning electron microscopy of HeLa cells expressing RhoG^WT^ (a) or RhoG^G12E^ (b). Confocal images of phalloidin-stained (red) F-actin and DAPI-stained (blue) nuclei in RhoG^WT^ (c) or RhoG^G12E^ (d) cells. Scale bars, 20 µm. **(B)** Cell spread area (µm^2^) quantified from phalloidin-stained boundaries using Fiji (n = 30 cells per condition from 3 independent experiments). Data are mean ± s.e.m. Dots represent individual cells. ****P < 0.0001 (Statistical significance was determined by Mann-Whitney test).

### G12E mutant enhances focal adhesion formation and decreases cell migration

Given the increased cell–substrate contact area and cytoskeletal reorganization, we examined potential changes in focal adhesions of cells expressing the RhoG^G12E^ variant during collective migration induced by wound healing. The focal adhesions (FAs) were visualized with an anti-vinculin antibody (**Fig. 6A**). Our results revealed that cells expressing RhoG^G12E^ exhibited a significant increase in the number of focal adhesions per µm^2^ compared with both RhoG^WT^ expressing cells and parental cells (**Fig. 6B and Supp. Fig. 5A**). In addition, the average size of individual focal adhesions was markedly larger in RhoG^G12E^-expressing cells (**Fig. 6C**), suggesting enhanced FA maturation. Similar results were obtained in MCF10A epithelial cells stably expressing RhoG^G12E^ or RhoG^WT^ (**Supp. Fig. 5B**).

**Figure 6:**
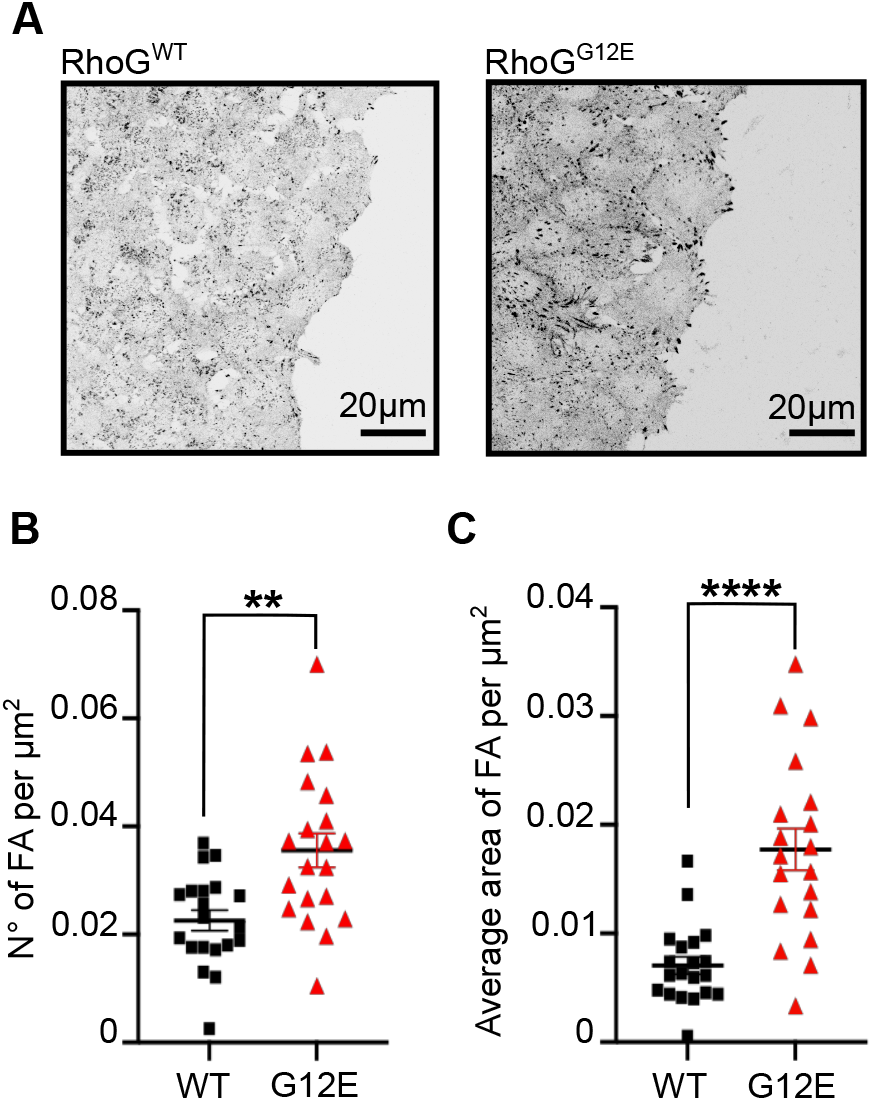
RhoG^G12E^ expression leads to an increase in focal adhesions number and size. RhoG^G12E^ expression leads to an increase of focal adhesions in number and in size. **(A)** Confocal Z-stack images of migrating RhoG^WT^ or RhoG^G12E^ cells toward a wound, stained for vinculin (focal adhesions). Scale bars, 20 µm. **(B)** Focal adhesion density (number/µm^2^) and **(C)** average focal adhesion area (per µm^2^) per cell. n = 20 cells per condition from 3 independent experiments. Data are mean ± s.e.m. Dots represent individual cells. **P < 0.01, ****P < 0.0001 (Statistical significance was determined by Mann-Whitney test).

Finally, given the increased focal adhesions of RhoG^G12E^ mutant expressing cells, we assessed collective cell migration by performing wound-healing assays. As RhoG^G12E^ may exhibit a slight decrease in cell proliferation rate (**Supp. Fig. 6A and B**), the migration assays were performed in the presence of mitomycin C to avoid confounding effects of proliferation. (**Movies 1-3**). At 30 hours, RhoG^WT^ cells had nearly closed the wound, whereas substantial gaps remained in monolayers expressing RhoG^G12E^. Using FiJi, centroid movements of cells located at the leading edge were determined at 24 hours **(Fig. 7B)** and used to determine the migration distance of the cells **(Fig. 7C)**. Quantitative analysis of the migration distances of cells located at the leading edge of the wounds revealed that RhoG^G12E^-expressing cells migrated significantly slower than RhoG^WT^-expressing cells. By 24 hours, the difference in migration reached almost a twofold difference (**Fig. 7A and C**).

**Figure 7:**
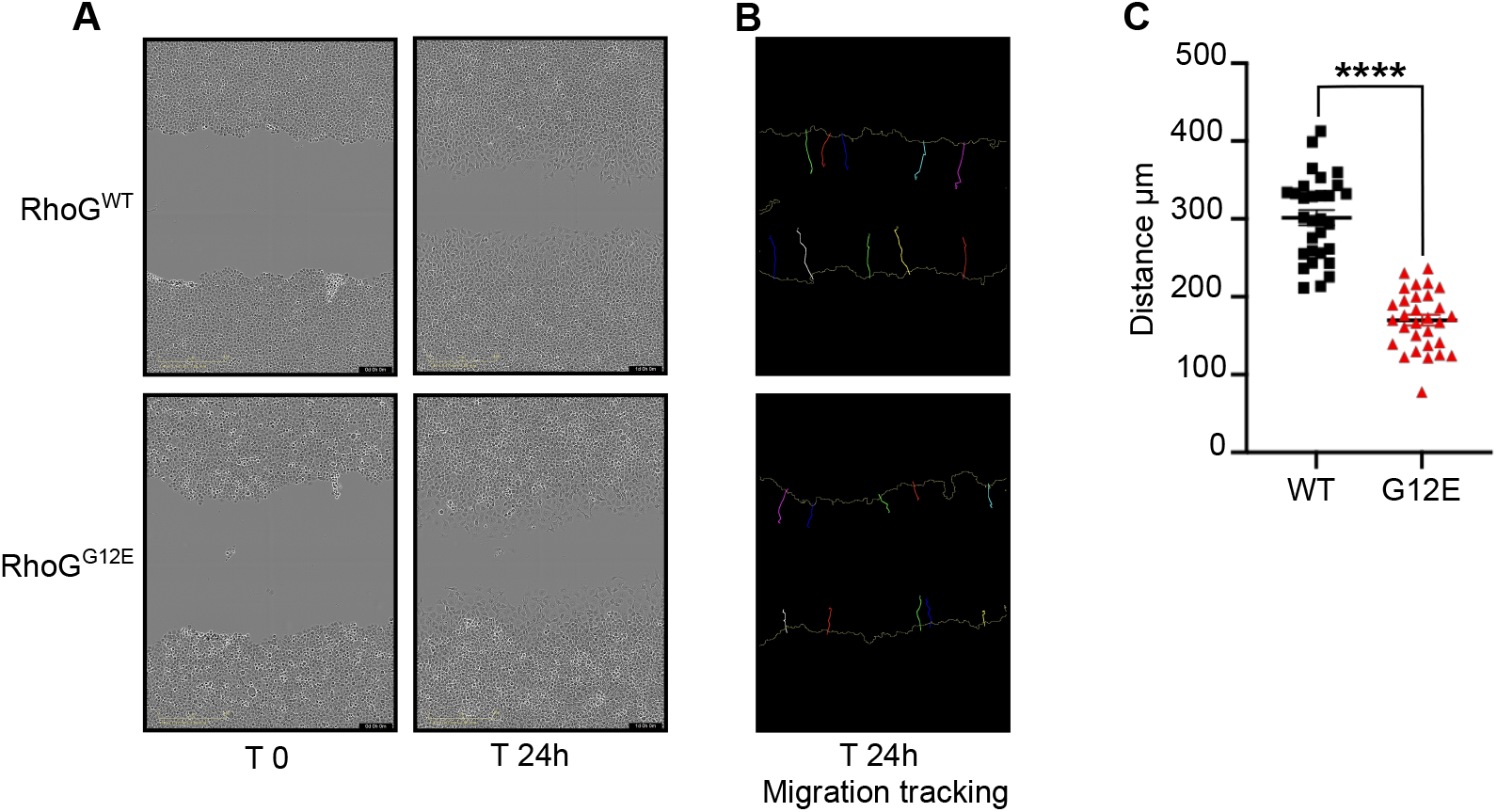
RhoG^G12E^ expression decreases migration rate in collective migration assays. **(A)** Incucyte images of scratch-wound closure by HeLa RhoG^WT^ or RhoG^G12E^ cells at 0 h and 24 h. **(B)** Trajectories of leading-edge cells over 24 h determined from centroid movement. **(C)** Migration distance (µm) toward the wound (n = 30 cells per condition from 3 independent experiments). Data are mean ± s.e.m. Dots represent individual cells. ****P < 0.0001 (Statistical significance was determined by Mann-Whitney test).

## Discussion

### RhoG^G12E^ phenocopies RhoG-depletion by breaking the cycle

Rho GTPases signaling relies on continuous nucleotide cycling [16],[17],[18] rendering the physiological impact of mutations more difficult to predict. Yet Ras-derived “constitutively active” mutants remain the default tools used to probe Rho function [19],[20]. While nearly all G12 variants blunt intrinsic GTP hydrolysis, this characteristic has become a poor discriminator of their functional differences in cells. The meaningful distinctions lie elsewhere: they arise from how each substitution redistributes the balance between GEF-driven activation and GAP-mediated inactivation. We show that hydrolysis is strongly impaired in RhoG^G12E^, but also that nucleotide exchange is not fully preserved. GDP and GTP release are both slowed, without a measurable change in nucleotide affinity. Functionally, we show that the RhoG^G12E^ mutant does not bypass upstream activation inputs and a GEF input remains necessary. Although GEF-mediated exchange is reduced, the magnitude of this defect is modest compared to the dramatic loss of GAP responsiveness. For GAPs, the magnitude of the observed defect exceeds what would be expected from reduced intrinsic hydrolysis alone and instead implies a failure to form or stabilize the GAP-supported transition state. We expect two consequences: i) buildup of the GTP-bound pool of RhoG^G12E^ upon an activation signal is slower than for Ras G12 mutants with an intact nucleotide exchange rate, because their GEF-mediated activation rate remains limited, ii) once molecules do reach the active state, they stall since we show that GAP-dependent clearance is ineffective. Consequently, the shift toward GTP-bound RhoG^G12E^ is probably not abrupt but cumulative, building over time as activation events are no longer efficiently erased. Consistent with this hypothesis, we show that early migration dynamics remain comparable between RhoG^WT^ and RhoG^G12E^. The delayed divergence in migration kinetics is consistent with progressive imbalance in regulatory cycling **(Movies 1-3)**.

In *in vitro* assays we show that ELMO1 binding is unaffected when the nucleotide-bound state is controlled, indicating that the mutation does not alter effector recognition. In cells, the increased effector association reflects progressive accumulation of active RhoG^G12E^ and not altered effector binding. Although we did not directly compare RhoG^G12E^ expression with RhoG depletion, the observed phenotypes resemble previously reported RhoG knockdown effects [10],[11]. At the functional level, we show that RhoG^G12E^ expression is accompanied by enhanced spreading, increased focal adhesions and reduced migration, all of these cellular phenotypes being consistent with the decrease of RhoG signaling [10],[11]. This behavior fits a broader principle proposed in multiple studies of Rho-family GTPases: locking the switch in the “ON” position is not equivalent to keeping the pathway active [4],[5]. Models of Rho GTPase dynamics, as well as experimental perturbations that dampen cycling without abolishing activation, converge on the same point: persistence without turnover degrades signal transmission over time [21](**Figure 8**).

**Figure 8:**
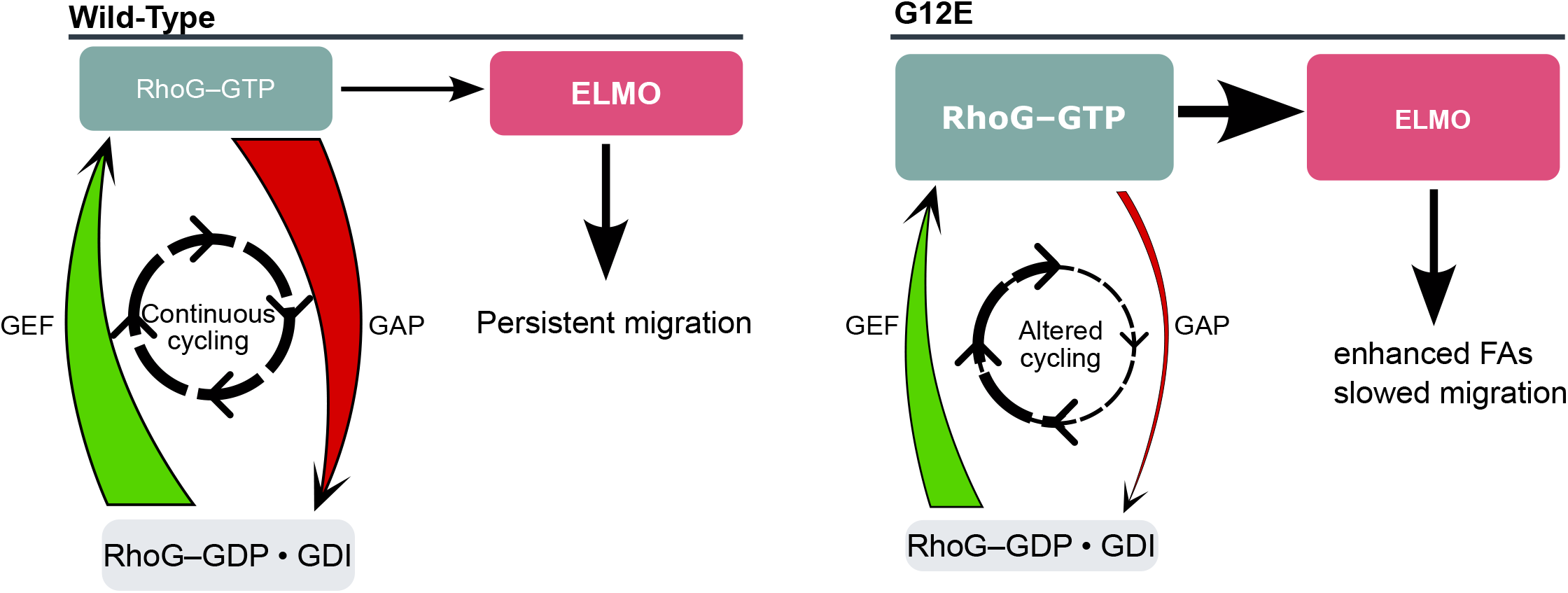
Summary of the RhoG^G12E^ biochemical profile and cellular output Scheme of the proposed mechanism. Across the cycle, G12E produces an asymmetric kinetic phenotype: activation by GEFs is reduced but remains functional, GTP intrinsic hydrolysis is strongly impaired, and GAP-catalyzed inactivation is nearly abolished. This combination predicts a GTP-bound population that can be generated but cannot be efficiently cleared, setting the stage for persistence without productive turnover in cells. Wild-Type RhoG displays a continuous nucleotide cycling through GEF and GAP activities. On the other hand, G12E mutant, due to its abolished hydrolysis activity does not cycle and tend to accumulate, with time, in an active form. This altered cycling leads to slowed of migration and enhanced Focal adhesion.

Seen from this angle, RhoG^G12E^ is not a constitutively active mutant in the usual sense. Its phenotype reflects a functional dysregulation of RhoG-dependent processes placing it closer to loss-of-function than gain-of-function. That inversion is easy to miss if one equates GTP loading with activity. In a pathological context, this inversion is critical: mutations such as RhoG^G12E^ may drive disease not by hyperactivating signaling pathways, but by inhibiting the dynamics of the GTPase cycle required for coordinated migration, polarity, and tissue organization. Yet RhoG^G12E^ clinical significance remains uncertain, and our study does not establish causality. ClinVar identified the mutation in a patient with an immune disorder. Thus, it is tempting to compare the G12E variant to a loss of RhoG function in a disease context. Recent work has shown that RhoG deficiency due to compound heterozygous mutations in the RHOG gene leads to HLH and altered cytotoxic function in T lymphocytes [22].

### A note on using hydrolysis-defective mutants as functional proxies

Altogether, these observations argue for caution when deploying RhoG mutants at position 12 as a “constitutively active” tool in cellular assays. Our data instead support using this mutant as a functional proxy for RhoG loss, at least in contexts where signaling output depends on sustained cycling. This raises a broader, and largely unresolved, issue in the Rho field. In contrast to the classical interpretation of Ras, where Gly12 mutations have been biochemically dissected in enough detail to justify their routine use as constitutively active alleles, equivalent assumptions for Rho-family GTPases remain weakly grounded. Hydrolysis-defective mutations are often treated as interchangeable gain-of-function reagents, yet systematic biochemical evaluation remains limited. These findings suggest that Gly12 substitutions in Rho-family GTPases may not uniformly behave as constitutively active alleles and warrant careful biochemical evaluation. Our results suggest that without a detailed understanding of biochemical and functional consequences of a mutation, interpretation of cellular phenotype is ambiguous. To our knowledge, it remains unclear whether any Rho GTPase Gly12 mutant genuinely behaves like Ras Gly12, specifically its ability to constitutively induce downstream signaling without disrupting it. The answer to this question remains unclear and warrants further investigation in future studies.

## Materials and Methods

### DNA cloning and sequences

For bacterial expression and production, DNA sequences encoding amino acids 1–167 of RhoG wild-type and of the G12E mutant were synthesized, codon-optimized, and cloned into the pET28 expression vector by Twist Bioscience.

For mammalian expression, the human cDNA sequence of RhoG (Genbank accession number NM_001665) was obtained from Origene Human lentiOrf library, subcloned and inserted into pLex lentiviral expression plasmid (Thermo Scientific) by SLIC assembly at the Cellule d’Ingénierie Génétique et d’Expression (CIGeX-CEA-Paris-Saclay / IBFJ-iRCM; Fontenay aux roses; France). The G12E mutant was obtained by single nucleotide mutagenesis. All sequences were confirmed by DNA sequencing.

### Bacterial protein expression, and purification

RhoG and the DHPH domain of TRIO were expressed and purified as previously described (Peurois *et al*., 2017). Constructs encompassing the C1GAP domain of RacGAP1, the GAP domain of BCR, and the DHPH domain of PREX1 were expressed in *E. coli* BL21(DE3) Star strain (ThermoFisher Scientific) grown in Terrific Broth (from ThermoFisher Scientific). Protein expression was induced at OD_600_ ≈ 0.6 with 0.5 mM IPTG for 16 h at 20°C. Cells were resuspended in lysis buffer (50 mM HEPES pH 8.0, 300 mM NaCl, 30 mM imidazole pH 8.0, 5% glycerol, 2 mM β-mercaptoethanol) supplemented with protease inhibitors and benzonase, and lysed by sonication. Clarified lysates were purified by Ni-NTA affinity chromatography (Cytiva) and eluted with lysis buffer containing 300 mM imidazole. Proteins were further purified by size-exclusion chromatography on a Superdex 75 column (Cytiva) equilibrated in 50 mM HEPES pH 8.0, 150 mM NaCl, 5% glycerol, and 2 mM DTT, concentrated, flash-frozen, and stored at −80 °C.

### GAP and GEF activity assays

RhoG wild-type and RhoG^G12E^ were preloaded with nucleotides as previously described [23]. Briefly, 100 µM RhoG was incubated with 10 mM EDTA and 250 µM BODIPY-FL-GDP or BODIPY-FL-GTP (Jena Bioscience) for 30 min at room temperature. Loading was stabilized by addition of 75 mM of MgCl_2_, and excess nucleotide was removed by desalting using PD SpinTrap G-25 columns (Cytiva). EDTA-stimulated exchange were performed with 20mM of EDTA. GEF-mediated nucleotide exchange was monitored at 25 °C by the decrease in BODIPY fluorescence (excitation 485 nm, emission 530 nm) using a FLEXstation 3 microplate reader (Molecular Devices). Reactions were performed in HKM buffer (50 mM HEPES pH 7.4, 120 mM potassium acetate, 1 mM MgCl_2_) with 1 µM RhoG. TRIO DHPH and PREX1 DHPH were used at concentrations ranging from 1–150 nM and 1–100 nM, respectively. Where indicated, EDTA was added to a final concentration of 5 mM at the start of the measurement.

GAP assays were performed with 1 µM GTP-loaded RhoG in the presence of 10 µM MDCC-phosphate-binding protein as described previously [14],[24]. C1GAP RacGAP1 and BCR GAP domains were used at concentrations ranging from 20–200 nM.

For both GEF and GAP assays, observed rate constants (k_obs_) were obtained by fitting fluorescence traces to a mono-exponential function under pseudo-first-order conditions. Apparent catalytic efficiencies (k_cat_/K_M_) were determined from linear regression of k_obs_ as a function of GEF or GAP

### Cell culture and generation of stable cell lines

HeLa (ATCC # CRM-CCL2) cells, stable HeLa cell lines expressing exogenous RhoG^WT^ or RhoG^G12E^ variants and 293T cells (ATCC# CRL-3216) were cultured in Dulbecco’s modified Eagle medium (DMEM) (Gibco, 31966-021) supplemented with 10% (v/v) FBS (Eurobio Scientific, CVFSVF00-01) and 1% (v/v) penicillin/streptomycin at 37°C in a humidified incubator with 5% CO_2_. Lentiviral particles were produced by calcium phosphate co-transfection of 293T cells with pGAG, pEnv-VSVG expression vectors and plasmid encoding a self-inactivating lentivirus vector containing the inserts of interest. Culture supernatants were harvested 72 hours post-transfection and were centrifuged three times at 3000 g. Viral supernatants were stored at −80°C. Viral transductions were conducted by adding one volume of viral supernatant of interest to one volume of the indicated cell culture medium. Culture medium was replaced after 18 hours of transduction by fresh complete medium. Depending on the conditions, cells underwent or not selection 24 hours post-transduction. Stable cell lines were selected by addition of 1µg/mL puromycin (ThermoFisher Scientific, J67236.XF) to the culture medium.

### Immunoblotting

Before lysis, cells were washed with ice-cold PBS and lysed on ice for 30 min in RIPA buffer (20 mM Tris pH 8, 150 mM NaCl, 10 mM Na_2_HPO4, 5 mM EDTA, 1% NP40, 1% DOC, 0.1% SDS, and 10% Glycerol) supplemented with protease and phosphatase inhibitors (Thermo Fisher Scientific, A32961) and 1 mM sodium orthovanadate. Post nuclear supernatants were recovered after 20 minutes centrifugation at 11,000 g and 4°C. Protein concentrations were determined using a BCA Protein Assay (Thermo Fisher Scientific, 23225) and proteins were denatured in reducing Laemmli’s sample buffer (Sigma, S3401-1VL) at 95°C for 5 min for subsequent western blot analysis. Proteins were resolved by SDS–PAGE, transferred onto 0.2 µm PVDF membranes (Cytiva) and detected using the Clarity Western ECL Substrate (Bio-Rad, 170-5061) and a ChemiDoc imaging system (Bio-Rad). Band intensities were quantified using Fiji and normalized to GAPDH signal. Experiments were performed using three independent experiments. Antibodies and dilutions were as follows: anti-RhoG (BioLegend, 814701, 1:2000), anti-GAPDH (Santa Cruz, sc-166545, 1:1000), and HRP-conjugated anti-mouse secondary antibody (Sigma, NA931V, 1:5000).

### RhoG activation assay (ELMO1 pull-down)

GST-tagged ELMO1 was expressed in *E. coli* and purified as described in (SM Goicoechea et al; J. Cell Science; 2017). Purified GST–ELMO1(a kind gift by R. Garcia-Mata; University of Toledo) was immobilized on glutathione-coated magnetic beads (Pierce, Thermo Fisher Scientific, Ref. 78601) by incubation at 4 °C for 1 hour under gentle rotation and store in activity buffer 50 mM HEPES pH 7.5, 100 mM NaCl, 10 mM MgCl_2_, 1% Triton X-100, 10% glycerol, 1 mM DTT, protease and phosphatase inhibitors (Thermo Fisher Scientific, A32961).

For *in vitro* pull-down assays, purified RhoG^WT^ or RhoG^G12E^ were preloaded with GDP or GTPγS (Jena Bioscience, 127646) and incubated with 10 µg GST–ELMO1 loaded on magnetic beads for 30 min at 4 °C. Beads were washed extensively, and bound proteins were eluted by boiling in SDS sample buffer. Eluted proteins were analyzed by SDS–PAGE followed by Coomassie Blue staining.

For cellular pull-down assays, HeLa cells from 10-cm tissue culture dishes and harvested near confluency were lysed in 0.5ml activity lysis buffer (50 mM HEPES pH 7.5, 100 mM NaCl, 10 mM MgCl_2_, 1% Triton X-100, 10% glycerol, 1 mM DTT, protease and phosphatase inhibitors). Lysates were clarified by centrifugation at 13,000 rpm for 10 min at 4°C and protein concentrations were measured using Protein Assay Kit. Equivalent amount of total protein were incubated with 10 µg of GST–ELMO1 loaded on magnetic beads for 30 min at 4°C under gentle rotation, washed three times with activity buffer, and bound proteins were eluted and analyzed by SDS–PAGE followed by immunoblotting for RhoG as described above.

### Cell size quantification

For cell size quantification, under confluent HeLa cells grown on coverslips were fixed at room temperature for 30 min in 4% paraformaldehyde (PFA; EM grade, CliniSciences Ref#15710), blocked in PBS, with 1.5% BSA for 1h, incubated in permeabilization buffer (PBS-1X, 1.5% BSA, 0.5% Triton X-100) for 1h at room temperature. Afterwards, cells were stained with Alexa FluorTM 488 phalloidin conjugate (ThermoFisher Scientific, A12379) diluted in permeabilization buffer (1:500) for 1h. Nuclei were counterstained using DAPI (4’,6-diamidino-2-phenylindole, ThermoFisher Scientific, 62248) and coverslips were mounted in EprediaTM PermafluorTM mounting medium (Fisher Scientific). Confocal imaging was performed on a Leica TCS SP8 (Leica microsystems) equipped with an oil 63x (N.A. 1.32) objective. Cell surface size was measured from confocal z-stacks projections of Phalloidin staining by manually drawing polygonal selections around individual cells in maximum intensity projections using Fiji. Statistical quantification was performed on 30 cells per condition.

### Scanning Electron Microscopy (SEM)

HeLa cells were grown on Lab-Tek II slides (Nunc) in complete medium, washed three times with cold 0.1M NaPO_4_ buffer pH7.4 and fixed overnight at 4°C in Karnovsky’s fixative solution (2% paraformaldehyde and 2.5% glutaraldehyde). After one phosphate buffer wash, the cells were subjected to gradient dehydration by means of successive 3-minute incubations in ethanol solutions at 30%, 50%, 70%, twice at 80%, 90% and 100% then air-dried under a hood for 30 min. The samples were sputter-coated (Emitech K650) with a thin layer of gold (about 10 nm) for 4 minutes with sputter current of 60 mA. Images were acquired using a Hitachi S-3400N Scanning Electron Microscope.

### Focal adhesion assay

Cells were grown on glass coverslips as confluent monolayer, serum starved in DMEM medium containing 0.5% FBS for 1h prior being wounded by scraping with a tip. Non-adherent cells were removed by one wash in DMEM 0.5% FBS and complete medium was gently added. Cells were further incubated at 37°C with 5% CO_2_ for 5h. Slide preparation was performed according to the Actin Cytoskeleton / Focal Adhesion Staining Kit protocol (Sigma, FAK100). Briefly, cells were fixed with 4% PFA in PBS for 20 min at room temperature, followed by two washes with wash buffer (PBS containing 0.05% Tween^®^20). Blocking was performed for 30 min at room temperature with 1% BSA in PBS. Cells were incubated with mouse anti-Vinculin (1:100) in blocking solution for 1 hour at room temperature, washed three times with wash buffer, then incubated with Alexa Fluor™ 488 secondary antibody (ThermoFisher Scientific, Ref#A-11001) and TRITC-phalloidin (Sigma, FAK100) for 45 minutes. After washing, nuclei were stained with DAPI, coverslips mounted on glass slides and imaging was performed as described for cell quantification. Focal adhesion density (area of focal adhesion/µm^2^ of cell) was calculated by normalizing summed adhesion area to cell area, as described in [25] [26]. Quantification was performed on 20 cells per condition across three independent experiments.

### Wound healing assay

Wound healing assay and live cell imaging was carried out with Incucyte® Live-Cell Analysis System. Confluent cells plated on Incucyte® Imagelock 96-well plate (Sartorius BA-04588) were pre-treated for 1h with 10µg/mL Mitomycin C (Sigma M5353) and serum-starved in DMEM medium containing 0.01% FBS for 1h. Scratch was created with Incucyte® Woundmaker followed by immediate gentle washes to remove non-adherent cells. Phase-contrast images were acquired every 3h for 48h at 37 °C under 5% CO2 with a 10x objective. Quantification of 30 cells migration per experiment was performed by manually tracking of nuclear trajectories of cells at the wound edge across all timepoints using Fiji. Movies were then compiled from these images in Fiji.

### Statistical analysis

All statistical analyses were performed in GraphPad Prism (version 10.4.2). Sample size (n) refers to independent experiments, and all quantifications were repeated on at least three independent experiments. Data are presented as mean ± s.e.m., unless otherwise stated. Groups were compared using either one-way ANOVA followed by adapted post hoc test, or Student’s t-test for pairwise comparisons. The exact P values, number of independent experiment (n), and type of statistical tests applied are indicated in figure legends.

## Acknowledgements

This study was supported by grants to A.N. from the French National Research Agency (ANR-22-CE11-0004) and to F.G. from AAP-Emergence-2022 (OI-HEALTHI-Université Paris-Saclay) and from AAP-Tecnics-2023 (GS-HEADS-Université Paris-Saclay). We are grateful to the scientific staff at Curie Orsay Imaging platform for their expert help and advice. We thank IDA ENS Paris-Saclay for their support. We thank R. Garcia-Mata (University of Toledo) for providing us with ELMO1 plasmid and Alexis Gautreau (BIOC, CNRS, École Polytechnique, Institut Polytechnique de Paris) for providing with MCF10a cells.

## Author contributions

S.W., Y.F., C.P., M.Z., R.G., F.G and A.N. performed the experimental studies. Y.F., A.N. and F.G. carried out the analysis. A.N. and F.G. supervised the work. A.N. and F.G. wrote the manuscript with the input from other authors.

