## Supplementary material for "Persistence without turnover: the RhoG G12E mutant highlights the role of nucleotide cycling in RhoG signaling": Supp Figures

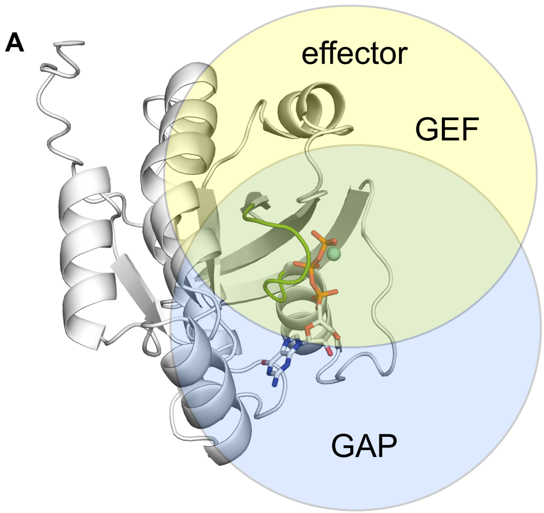


**Supplementary Figure 1. Model of RhoG G12E mutant. (A)** Structural model of RhoG G12E in complex with GTP predicted by AlphaFold3. The P-loop is highlighted in green, GTP and $Mg^{2+}$ are shown. Regions involved in GEF and effector interactions are indicated by a yellow circle, whereas surfaces implicated in GAP binding are highlighted in blue. **(B)** Close-up view of the nucleotide-binding pocket. The P-loop residue Gly12 is substituted by Glu12, introducing a negatively charged side chain in a confined region of the nucleotide-binding site. The $Mg^{2+}$ -coordinating residues Thr17 and Thr35 are shown. In the model, the Glu12 side chain is positioned near Gln61, a catalytic residue involved in the hydrolysis mechanism, suggesting that the substitution may perturb the geometry of the catalytic site and interfere with GAP and/or-GEF mediated regulation.

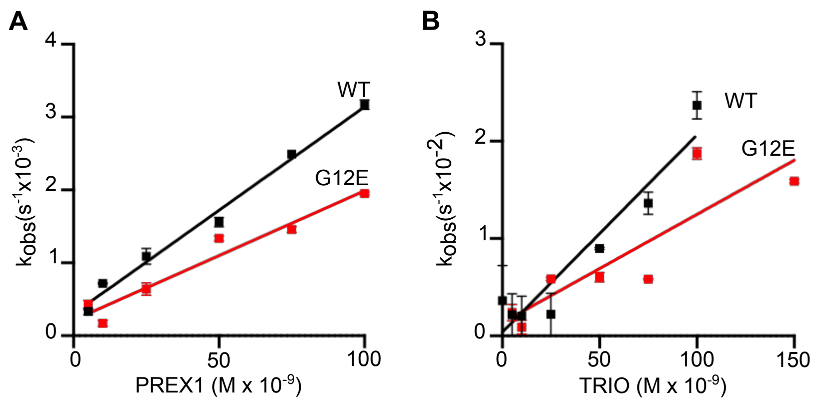


**Supplementary Figure 2.** The catalytic efficiency of **(A)** PREX1 and **(B)** Trio DHPH constructs was determined using fluorescence kinetics in solution. $k_{cat}/K_M$ values were determined from $k_{obs}$ obtained over a range of active sites concentrations (0 to 100 nM) from 3 independent experiments.

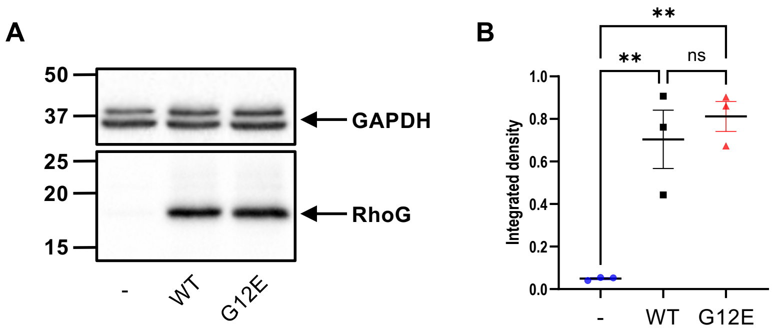


**Supplementary Figure 3: RhoG expression and quantification in HeLa cells. (A)** Expression of endogenous RhoG (-), lentiviral mediated overexpression of wild-type RhoG (WT) and mutant RhoG (G12E). GAPDH is used as a loading control. **(B)** Quantification of RhoG expression was performed by measuring integrated density of western blot bands using Fiji, normalized to GAPDH. Data represent mean ± standard error from independent experiments (n=3). Statistical significance was assessed by ordinary one-way ANOVA. $p < 0.01$ (**) denotes significant differences, ns = not significant.

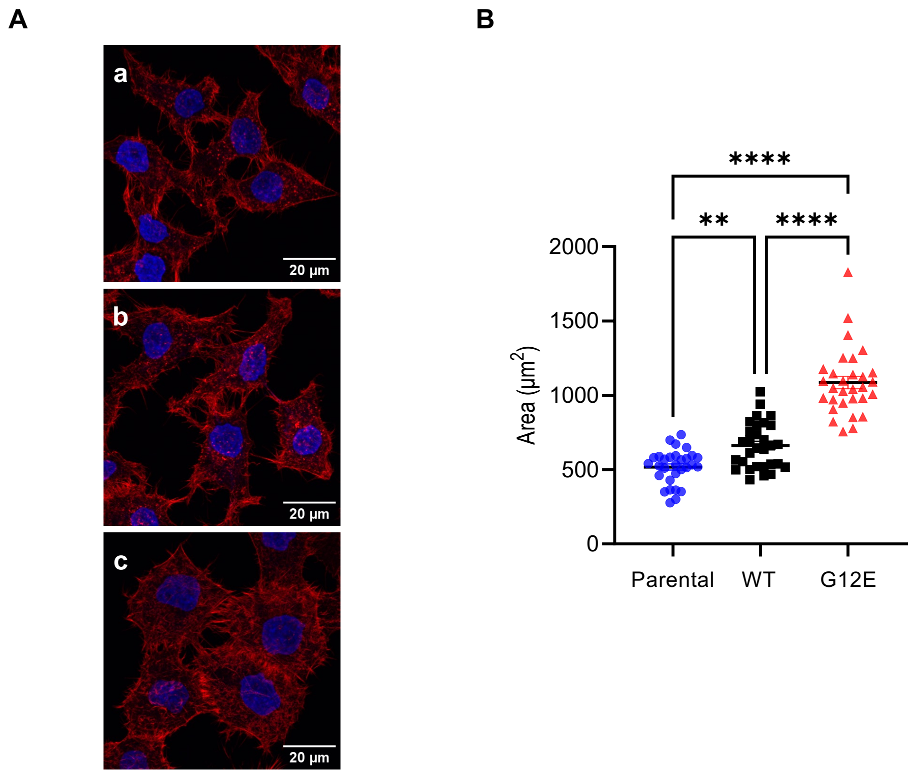


**Supplementary Figure 4: RhoG$^{G12E}$ increased cell spreading in HeLa cells. (A)** Confocal z-stack images depicting F-actin cytoskeleton in HeLa **(a)**, RhoG$^{WT}$ **(b)**, and RhoG$^{G12E}$ **(c)** cells stained with phalloidin (red fluorescence), and DAPI (blue). **(B)** Quantification of cell spreading area (µm$^2$) of 30 cells each type using Fiji. Points show individual cell measurements; bars indicate mean ± standard error, independent experiment was repeated at least three times. Statistical analysis was conducted using ordinary one-way ANOVA. ** $p < 0.01$; **** $p < 0.0001$ denotes significant differences.

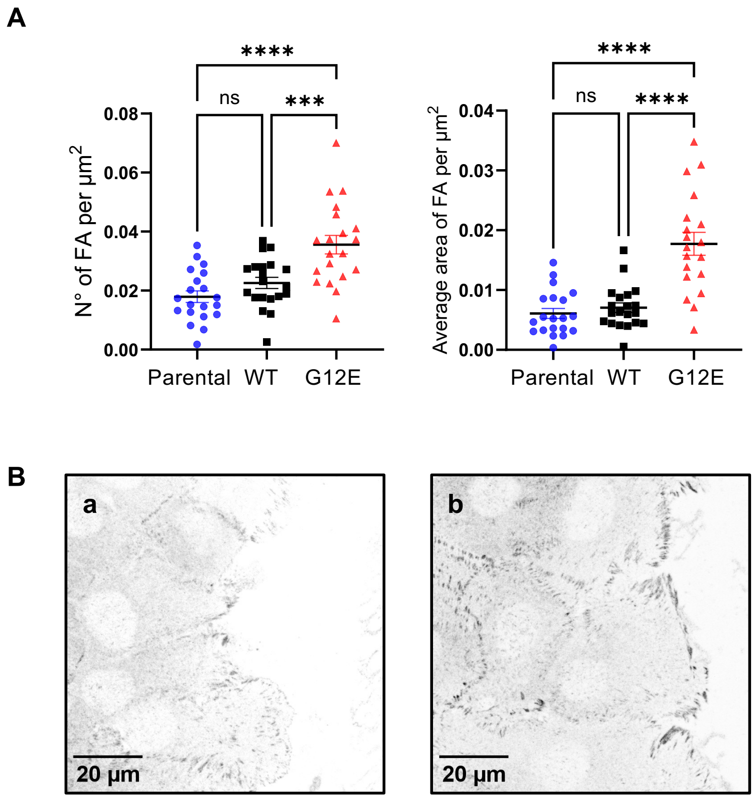


**Supplementary Figure 5: RhoG$^{G12E}$ increases focal adhesion number and size. (A)** Quantification of focal adhesion (FA) number and size per µm$^2$ of HeLa cells based on confocal images stained with vinculin. Data represent mean ± standard error of 20 cells per condition, independent experiment was repeated at least three times. Statistical analysis was conducted using ordinary one-way ANOVA. *** $p < 0.001$, **** $p < 0.0001$; ns, not significant. **(B)** MCF10a cells expressing RhoG$^{WT}$ **(a)** or RhoG$^{G12E}$ **(b)** stained with anti-vinculin.

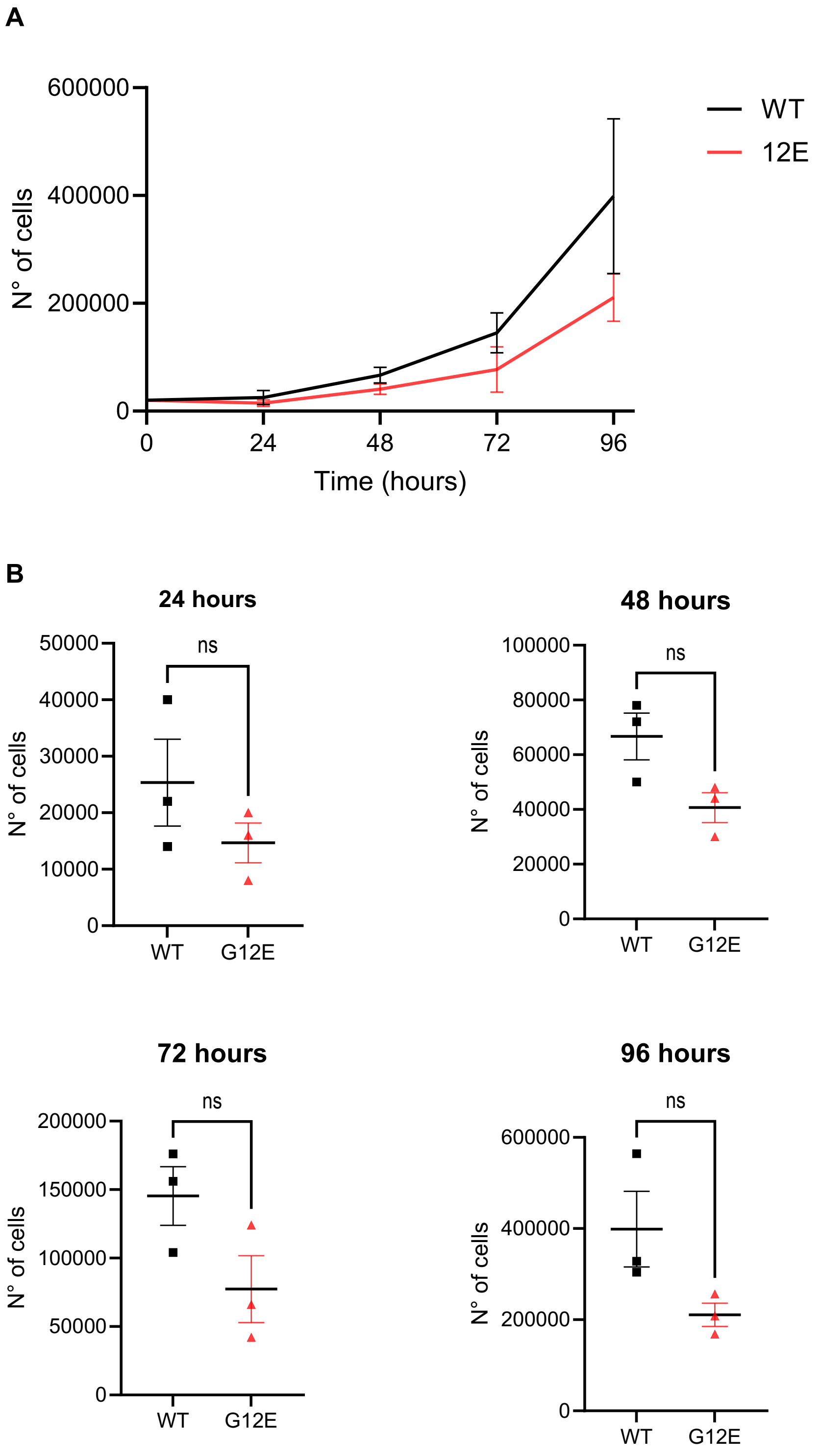


**Supplementary Figure 6: RhoG$^{G12E}$ expression does not substantially alter HeLa cell proliferation. (A)** Cell growth curves of RhoG$^{WT}$ and RhoG$^{G12E}$ HeLa cells over 96 h, quantified by manual counting every 24 h (n = 3 independent experiments). **(B)** Cell numbers at 24, 48, 72 and 96 h for RhoG$^{WT}$ and RhoG$^{G12E}$ cells. Data are mean ± s.e.m. from 3 independent experiments; dots represent individual experiments. ns, not significant (Statistical significance was determine by Mann–Whitney test).
